# Structural Brain Correlates of Speech Disfluency in Early Childhood: A Dimensional Analysis in a Non-Clinical Cohort

**DOI:** 10.1101/2025.08.18.670817

**Authors:** Ashmeet Jolly, Aura Yli-Savola, Elmo P. Pulli, Essi Saloranta, Henry Railo, Harri Merisaari, Ekaterina Saukko, Eero Silver, Venla Kumpulainen, Anni Copeland, Hasse Karlsson, Linnea Karlsson, Niina Junttila, Elina Mainela-Arnold, Jetro J. Tuulari

**Author notes:** Correspondence, FinnBrain Birth Cohort Study, Turku Brain and Mind Center, Kiinamyllynkatu 10, Medisiina A Building, 20520, Turku, Finland.

## Abstract

Most neuroimaging studies of speech disfluency have compared individuals who stutter with fluent controls. However, treating speech disfluency as a continuous, dimensional trait offers new insights into the neural basis of fluency during early childhood. This study aimed to investigate whether naturally occurring variation in speech disfluency is associated with grey matter structure in a non-clinical, population-based sample of 5-year-old children. The study included 120 participants (65M, 55F) from the FinnBrain Birth Cohort study. Speech disfluency was evaluated as a continuous measure from audiovisual speech samples, with transcription and analysis conducted using the SALT software. Ambrose & Yairi (1999) classification system was used to categorize speech disfluencies into stuttering-like (SLD) and other disfluency types. T1-weighted images obtained through magnetic resonance imaging were analyzed using voxel-based morphometry (VBM) with the CAT12 toolbox and complemented by surface-based morphometry with FreeSurfer. Whole-brain statistical analysis was employed to examine the association between grey matter metrics and speech disfluency. We found that VBM-derived proportional grey matter volume in the left middle frontal gyrus, left posterior cerebellum, and right superior frontal gyrus was positively associated with speech disfluency, specifically SLD, in children (*p* <.001; *p*=.002; *p* <.001, FDR corrected). No significant associations were found for cortical thickness or surface area. Additionally, no notable sex differences were observed. Our findings suggest that speech disfluency in early childhood is linked to localized structural differences in regions supporting motor planning and cognitive control, without broader changes in cortical thickness or surface area. Importantly, similar brain regions have been implicated in studies comparing children who stutter to those who do not, suggesting that normal variation in disfluency captures meaningful neurobiological differences even in non-clinical populations. This supports the value of treating speech disfluency as a spectrum and underscores the importance of longitudinal, multimodal research to clarify how these structural features evolve and influence later fluency outcomes.

## Introduction

Fluency comprises the seamless flow, pace, and ease of speech production (Schmidt, 1992). A speech disfluency refers to interruptions, irregularities, or non-lexical sounds that disrupt the flow of otherwise fluent speech (ASHA, 2024). These may include “false starts,” where words or sentences are abruptly cut off; phrases or syllables that are repeated or restarted; and “fillers,” such as grunts or semi-articulated sounds like “uh,” “erm,” “um,” or “hmm.” Occasional disfluency is a natural part of speaking for everyone. Individuals who stutter often experience a higher frequency of disfluencies and specific types of disfluencies. These may include repeating parts of words (repetitions), lengthening a sound for an extended duration (prolongations), or struggling to begin a word (blocks) (ASHA, 2024).

Stuttering-Like Disfluencies (SLD) and Other Disfluencies (OD) are two primary classifications that have been instrumental in both research and clinical settings to diagnose stuttering. In a landmark study which focused on stuttering in children aged 2–5 years (Ambrose & Yairi, 1999), these distinctions were defined based on the nature and frequency of disfluencies (Table 1). Subsequent studies reinforced the utility of this classification, with the criterion of ≥3% SLD *per 100 words* emerging as a viable standard threshold for distinguishing between children who stutter and children who do not stutter (Conture, 2001; Yairi & Ambrose, 2005). This criterion has demonstrated high sensitivity and specificity across several languages, including Dutch, German, Spanish, and English (Ambrose & Yairi, 1999; Boey et al., 2007; Carlo & Watson, 2003; Natke et al., 2006). While ODs often occur in fluent speech, they may also appear in the speech of people who stutter, albeit with less frequency and prominence compared to SLDs (Tumanova et al., 2014).

**Table 1.** Classification of speech disfluencies (Ambrose & Yairi, 1999; Jansson-Verkasalo et al., 2021)

| **Stuttering-Like Disfluencies** |  |  |
| --- | --- | --- |
|  | **Types** | **Description** |
|  | Part word repetition | Repetition of a sound or syllable within or in the beginning of the word |
|  | Monosyllabic word repetition | Repetition of a monosyllabic word |
|  | Prolongation | Prolongation of a sound resulting in an unusual duration of that sound |
|  | Block | Stopping airflow and sound during or before production of a vowel or consonant |
|  | Broken words | Stopping of airflow in the middle of a vowel within a word |
| **Other Disfluencies** |  |  |
|  | Filled Pause | Filler words or non-linguistic sounds within an utterance |
|  | Revision and abandoned sentences | Correction of word choice/grammatical or phonological errors/add or delete lexical information |
|  | Multisyllable word repetition | Repetition of a multi-syllabic word |
|  | Phrase repetition | Repetition of a phrase within an utterance |

Some researchers have argued that stuttering may not be a binary phenomenon but rather a gradient of speech fluency disruptions (Adams & Runyan, 1981) and this has been reinforced by recent multifactorial theories (Smith & Weber, 2017; Tichenor & Yaruss, 2021). Stuttering and disfluency may exist along a continuum, reflecting complex interactions between motor, cognitive, linguistic, and emotional factors. This highlights the importance of considering speech fluency as a spectrum rather than relying solely on binary classification. Fundamentally, even if stuttering ultimately reflects a categorical difference, where individuals who stutter exhibit brain abnormalities distinct from those underlying typical disfluencies, understanding the neural correlates of normal disfluency is essential for identifying what may be unique to clinical stuttering. Conversely, if speech disfluency lies on a true continuum, then the brain correlates with typical disfluencies should parallel those found in clinical stuttering. Thus, examining natural variation in speech disfluency provides critical insights for both theoretical models and clinical understanding of fluency disorders.

Studies examining structural differences in the brains of those who stutter have consistently reported altered structures in multiple brain regions, particularly in grey matter (Etchell et al., 2018). A systematic review of these studies (Table 2) highlights consistent regions implicated in stuttering, providing a benchmark for evaluating the brain-speech disfluency relationships in our sample. It is important to note that the studies summarized in Table 2 focus exclusively on structural grey matter and employ voxel-based morphometry (VBM); (Ashburner & Friston, 2000) or FreeSurfer (Fischl, 2012) for analysis. These methods are among the most widely used in neuroimaging to examine grey matter morphology. VBM allows for a voxel-wise comparison of grey matter density or proportional volume across participants, while FreeSurfer provides detailed measurements of cortical thickness and surface area based on anatomically defined regions-of-interest. By concentrating on these two methods, the table highlights studies that leverage these techniques to analyze structural variations in grey matter, offering a consistent basis for comparison across findings. All included studies utilized clinical samples (diagnosed persistent/recovered stuttering), as no previous structural neuroimaging studies have examined subclinical or continuous variation in speech fluency.

**Table 2.** Overview of Neuroimaging Studies (2000-2024) Investigating Structural Grey Matter Differences in Stuttering.

| Study | Technique | Age (in years) | Disfluency/Stuttering assessment method used | Key results |
| --- | --- | --- | --- | --- |
| (Jäncke et al., 2004) | VBM | 10 AWS<br>10 AWDS<br>(Mean age = NA) | SSI | No discernible disparity in GMV between AWS and AWDS. |
| (Beal et al., 2007) 8/18/25 2:12:00 PM | VBM | 26 AWS (Mean age = 30.29); 28 AWDS (Mean age = 30.53) | SSI | AWS showed greater proportional GMV compared to AWDS in the left inferior frontal gyrus as well as bilateral superior temporal gyrus. |
| (Chang et al., 2008) | VBM | 8 persistent CWS<br>7 recovered CWS<br>7 CWDS<br>(Mean age = 9-12) | (Ambrose & Yairi, 1999) criteria | Less GMV in the left middle frontal gyrus was found in recovered stutterers compared to persistent stutterers. |
| (Kell et al., 2009) | VBM | 13 persistent AWS (Mean age = 18-39); 13 recovered AWS (Mean age = 16-65); 13 AWDS (Mean age = 23-44) | (Boberg & Kully, 1994) criteria | AWS showed a decrease in GMV in the left inferior frontal gyrus which was negatively correlated with stuttering severity. |
| (Lu et al., 2010) 18.8.2025 14.12.00 | VBM | 10 AWS (Mean age = 19-31); 12 AWDS (Mean age = 22-29) | SSI-III | In comparison to AWDS, AWS showed more GMV in the left putamen, and less GMV in the left medial frontal gyrus as well as anterior superior temporal gyrus. |
| (Beal et al., 2013) | VBM | 11 CWS<br>11 CWDS<br>(Mean age = 6-12) | SSI-III | CWS had increased GMV in the right superior temporal gyrus and inferior frontal gyrus and had decreased GMV in the left inferior frontal gyrus and putamen in comparison to CWDS. |
| (Sowman et al., 2017) | VBM | 27 AWS (Mean age = 45.9)<br>27 AWDS (Mean age = 47.1) | Self-report | Smaller GMV in the left caudate nucleus was found in AWS relative to AWDS |
| (Garnett et al., 2018) | FreeSurfer | 11 recovered CWS<br>25 persistent CWS<br>34 CWDS<br>(Mean age = 3-11) | SSI-IV | Reduced cortical thickness was found in the left motor and premotor regions of those with persistent stuttering. |
| (Koenraads et al., 2019) | FreeSurfer | 26 CWS<br>489 CWDS<br>(Mean age = 6-9) | Parental reports | CWS showed thinner cortex in the left inferior frontal gyrus, and in bilateral frontal and parietal areas. |
| (Chow et al., 2023) | VBM | 23 recovered CWS<br>72 persistent CWS<br>95 CWDS<br>(Mean age = 3-12) | SSI-IV | CWS showed GMV reductions in speech-motor structures such as the putamen, nucleus accumbens, and thalamus, with distinct age-related shifts, as younger children primarily exhibit deficits in the putamen while older children show more pronounced anomalies in the thalamus and internal capsule. |

*Note: CWS – Children who stutter, CWDS – Children who don't stutter, AWS – Adults who stutter, AWDS – Adults who don't stutter, VBM – Voxel Based Morphometry, SSI – Stuttering Severity Instrument, GMV – Grey Matter Volume*

The table reveals that prior structural neuroimaging studies of stuttering have consistently identified structural differences involving regions crucial for speech and motor control. Frequently reported regions include the left inferior frontal gyrus, bilateral superior temporal gyrus, left middle frontal gyrus, left putamen, caudate nucleus, and motor/premotor cortical areas. Although exact directionality (increase or decrease in grey matter) varies, these regions are recurrently implicated in both child and adult studies, suggesting core neural circuits relevant to persistent stuttering.

From a neural perspective, (Theys et al., 2024) demonstrated that lesions causing acquired neurogenic stuttering converge on a network centered around the left putamen, claustrum, and amygdalostriatal transition area. The association of this lesion-based network with symptom severity in stuttering suggests a shared subcortical neuroanatomy underlying fluency disruption. While their primary focus was on subcortical structures, they also acknowledged potential contributions from cerebellar regions involved in motor control. These findings align with broader models of speech motor control e.g., (Guenther, 2016), which emphasize that fluent speech critically depends on the integrity of interconnected cortical, basal ganglia, thalamic, and cerebellar circuits. Drawing from this perspective, we propose that SLDs, reflecting disruptions in motor planning and execution, would be associated with structural differences within motor-cognitive loops involving the frontal cortex, basal ganglia, and cerebellum. Specifically, we suggest that even normal variation in the functioning of these circuits may contribute to individual differences in disfluency, providing insights into the full spectrum of speech fluency development.

In contrast, ODs which reflect linguistic processes such as lexical retrieval and linguistic monitoring, may correspond to variability in regions outlined in the dual-stream model (Hickok & Poeppel, 2007). According to this model, speech processing relies on dorsal (phonological-articulatory planning; left inferior frontal and parietal regions) and ventral streams (semantic-lexical processing; temporal cortices). Thus, individual differences in OD might specifically map onto these linguistic-processing cortical regions. Therefore, adopting a dimensional approach within a non-clinical, developmental population can provide novel insights into subtle neural variations underlying speech disfluency.

Surface-based analysis studies have shown varying results regarding cortical thickness and surface area in individuals who stutter. Although studies applying these methods to stuttering are limited, existing findings (Garnett et al., 2018; Koenraads et al., 2019) suggest cortical thickness differences particularly in frontal and motor cortices. Thus, examining both voxel-based and surface-based measures within the same population may yield complementary insights into neural correlates underlying speech disfluency.

Our study builds on previous neuroimaging research by incorporating established methodologies while introducing a novel dimensional approach to speech fluency, using continuous disfluency scores rather than categorical classifications. We conducted whole-brain analyses to examine the relationship between proportional grey matter volume (GMV) and disfluency, alongside complementary region-of-interest analyses focused on cortical thickness and surface area in 5-year-old children. Guided by theoretical models, we expected to find associations in the motor-cognitive loop for SLD and in regions supporting language processing for OD; however, given the exploratory nature of voxel-wise analyses, no specific hypotheses were made regarding the exact location or direction of effects.

## Methods

This study adhered to the principles outlined in the Declaration of Helsinki and received approval from the Joint Ethics Committee of the University of Turku and the Hospital District of Southwest Finland. Approval was granted under two protocols: (1) ETMK: 63/1801/2017 for speech assessments and (2) ETMK: 31/180/2011 for neuroimaging. The parents of the participants were informed about the visits and were requested to explain these to their child in a way that the child could understand. Consent from the child was also obtained.

The reporting was conducted following the STROBE (Strengthening the Reporting of Observational Studies in Epidemiology) guidelines, to ensure transparent and comprehensive reporting of our research.

### Participants

Participants were drawn from the prospective FinnBrain Birth Cohort study that began in 2010 and was established to study the effects of prenatal stress and early life stress on early childhood development from fetal life onwards. After birth, these children took part in longitudinal assessments entailing infant and early childhood brain and cognition (see Figure 1). An overview of the study and the population has been described previously (Karlsson et al., 2018). 120 (65 male, 55 female) Finnish speaking children successfully completed the speech assessment and structural neuroimaging visit suitable for the current study (please refer to Table 3 for demographic details).

**Figure 1.**
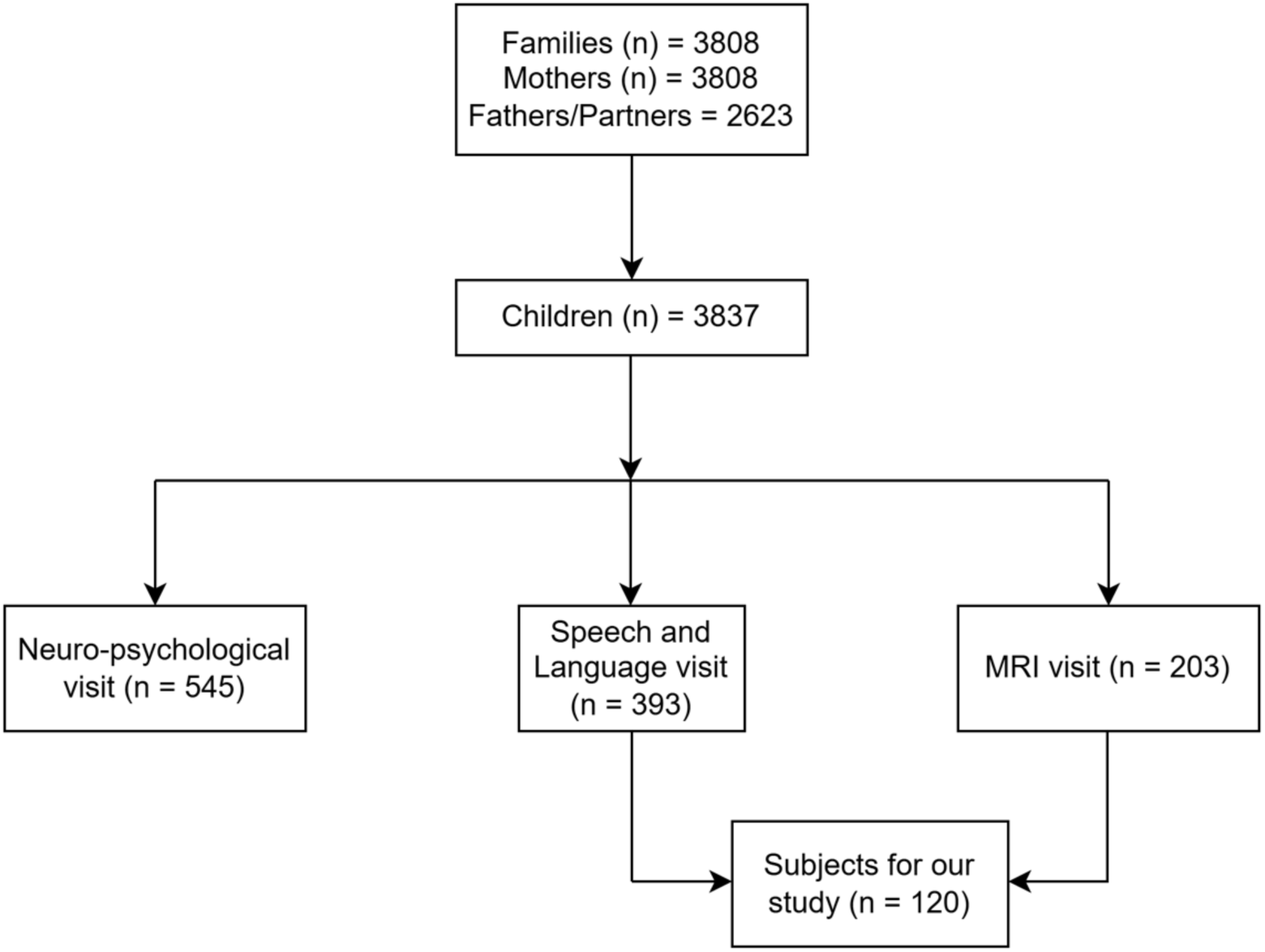
Flowchart of the participants in the current study (the number of children include 29 twin pairs)

**Table 3.** Demographics of the study population (n = 120)

| Measure | Mean | SD | Min | Max |
| --- | --- | --- | --- | --- |
| *Child Characteristics* |  |  |  |  |
| **Age at scan** (*in months*) |  |  |  |  |
| Boys | 65 | 1.7 | 62.8 | 69.4 |
| Girls | 64.9 | 1.3 | 63 | 69.1 |
| **Gestational Age at birth** (*in weeks*) |  |  |  |  |
| Boys | 39.7 | 1.7 | 33.9 | 42.3 |
| Girls | 40.1 | 1.3 | 36.3 | 42.3 |
| **Birth weight** (*in kg*) |  |  |  |  |
| Boys | 3.6 | 0.47 | 2.53 | 4.98 |
| Girls | 3.52 | 0.47 | 2.58 | 4.9 |
| **Head circumference at birth** (*in cm*) |  |  |  |  |
| Boys | 35.48 | 1.39 | 32 | 39 |
| Girls | 34.72 | 1.46 | 32 | 38.5 |
| **Handedness** |  |  |  |  |
|  | **Right-**<br>**Handed** | **Left-**<br>**Handed** | **Ambidextrous** | **Unknown** |
| Boys | 60 | 1 | 4 | 0 |
| Girls | 49 | 5 | 0 | 1 |
| *Maternal Characteristics* |  |  |  |  |
| **Age at birth** (*in years*) | 31 | 5 | 20 | 41 |
| **BMI at birth** (*in kg/m<sup>2</sup>*) | 24.17 | 4.19 | 17.48 | 41.95 |
| **Income** (*estimated after taxes, in euros*) | **Number** |  | **Percent** |  |
| ≤ 1500 | 33 |  | 27.5 |  |
| 1500-2500 | 66 |  | 55 |  |
| 2501-3500 | 13 |  | 10.8 |  |
| ≥ 3501 | 3 |  | 2.5 |  |
| Missing | 5 |  | 4.2 |  |
| **Education** |  |  |  |  |
| Low | 27 |  | 22.5 |  |
| Middle | 31 |  | 25.8 |  |
| High | 58 |  | 48.3 |  |
| Missing | 4 |  | 3.3 |  |
| **Background** |  |  |  |  |
| Finnish | 115 |  | 95.8 |  |
| Other | 1 |  | 0.8 |  |
| Missing | 4 |  | 3.3 |  |
| **Alcohol use during pregnancy** |  |  |  |  |
| Yes | 20 |  | 16.6 |  |
| No | 85 |  | 70.8 |  |
| Only before learning about pregnancy | 15 |  | 12.5 |  |
| **Smoking during pregnancy** |  |  |  |  |
| Yes | 3 |  | 2.5 |  |
| No | 113 |  | 94.1 |  |
| Only before learning about pregnancy | 4 |  | 3.3 |  |
| **Illicit drug use** (*gestational week 34*) |  |  |  |  |
| No | 117 |  | 97.5 |  |
| Missing | 3 |  | 2.5 |  |

*Note: BMI – Body Mass Index; Smoking status is based on FinnBrain questionnaires and National Institute for Health and Welfare (www.thl.fi); Maternal education, estimated monthly income, alcohol use were collected through questionnaires administered during the 14th gestational week.*

### Speech assessment visit

For the speech assessment of the 5-year-olds, we recruited families who had also participated in the neuropsychological study visits between 2017 and 2021. After contacting them, 393 children ended up participating in the study. Only Finnish speaking families were recruited for the speech and language study. The recruitment was done either via phone calls or emailing the families.

All children were monolingual Finnish speakers determined by asking the parent whether there were any other languages spoken in the family besides Finnish. Only children who heard or used Finnish 80% of the time or more participated.

From the initial pool of 394 children that were willing to participate in our study visit, 393 speech samples were acquired, as one child refused to speak. And out of 393, 325 speech samples were long enough (at least 300 words) for fluency analyses.

Speech samples were collected through audio and video recordings while the child interacted with the examiner in a free play setting. The examiners were speech-language pathology master’s students supervised by a clinical instructor. They spent approximately 20 minutes engaging in play with the child and encouraging them to speak by asking open-ended questions and making remarks about their play. Towards the end of the session, the examiner was instructed to create a bit of pressure on the child to increase the likelihood of disfluencies, by asking the child to quickly talk about an exciting event before leaving. The play sessions were audio and video recorded in a wave format using a Marantz recorder with a 44.1 kHz sampling frequency and 16-bit resolution.

### Magnetic Resonance Imaging Acquisition

The Siemens Magnetom Skyra fit 3T scanner with a 20-element head/neck matrix coil was utilized to scan the participants. The image acquisition process was accelerated using the Generalized Autocalibrating Partially Parallel Acquisition (GRAPPA) technique. As a component of a scan protocol lasting a maximum of 60 minutes, the MRI data was obtained. The current study obtained high resolution T1-weighted images with the following sequence parameters for its purposes: repetition time (TR) = 1,900 ms, inversion time (TI) = 900 ms, echo time (TE) = 3.26 ms, voxel size = 1.0 × 1.0 × 1.0 mm^3^, flip angle = 9 degrees, and field of view (FOV) 256 × 256 mm^2^. The scans were planned as per recommendations of the FreeSurfer developers (FreeSurfer Morphometry Protocols, 2009).

### MRI Visit

The target age for the neuroimaging visit was 63-65 months, and due to COVID issues there was a delay of 4-5 months in some of the cases (Pulli et al., 2022). Participants who met any of the following criteria were excluded from the MRI: 1) Participants born before gestational week 32, 2) Participants with abnormalities in development for example in senses or communicative abilities (e.g., deafness, blindness and congenital heart disease), 3) Participants who had an established long-term medical diagnosis (e.g., autism or epilepsy), 4) Participants with ongoing medical assessments (for example, a child was referred from a primary care setting to special healthcare), 5) Participants who use medication daily and continuously (e.g., oral medications, inhalants and topical creams. An exception to this was desmopressin (®Minirin)), 6) Participants with a history of head trauma, 7) Gold-plated metallic ear tubes (to assure good-quality scans), and routine MRI contraindications.

The scans were done by a team of four Ph.D. students, a research nurse and two radiographers. The participants were allowed to be awake or asleep during the scans, and a member of the research staff and a parent or guardian were present in the room throughout the entire scan. The participants were provided with earplugs and headphones, and they could watch a movie or TV show of their choice through mirrors that were placed inside the head coil. Foam padding was utilized to ensure the participants’ head stability and comfort during the scan.

Of the 203 children who participated in this visit, 146 completed both the neuropsychological and speech-language assessments. Among these 146, 8 did not undergo MRI scanning and 4 were excluded due to excessive motion artifacts in their T1-weighted images, resulting in 134 participants with usable MRI data. After MRI quality control (Pulli et al., 2022), 121 T1-weighted images remained. When these MRI datasets were merged with the speech assessment data, one participant was excluded due to missing or poor-quality speech data. Therefore, the final sample for the present study included 120 children with both high-quality MRI and speech-language data.

### Data Pre-processing

#### Speech Disfluency

The audio recordings of speech were transcribed using the SALT software v20.0 (Systematic Analysis of Language Transcripts) (Miller & Iglesias, 2020) using high-quality headphones. The final 300-350 words in each sample were examined for fluency. Words that were unintelligible or stand-alone affirmations or negations (e.g., yes, no, okay) were not included in the analysis, but those that were immediately followed by another word or phrase were retained (Ambrose & Yairi, 1999).

Disfluencies were then identified and categorized based on a system of disfluency types developed by (Ambrose & Yairi, 1999) and (Jansson-Verkasalo et al., 2021), with further details provided in Table 4 and Table 5.

**Table 4.**
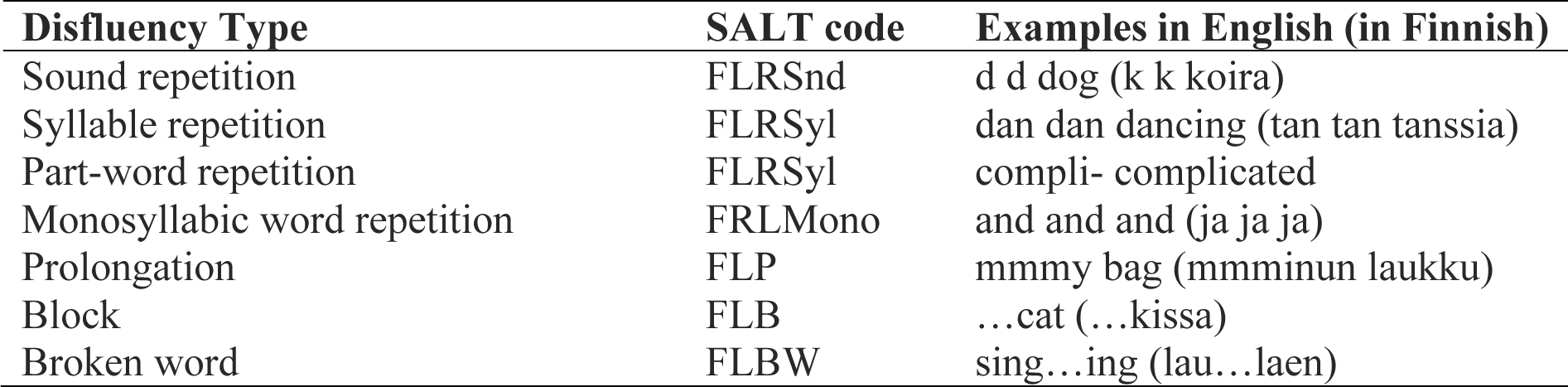
Stuttering-Like Disfluencies (SLDs)

| **Disfluency Type** | **SALT code** | **Examples in English (in Finnish)** |
| --- | --- | --- |
| Sound repetition | FLRSnd | d d dog (k k koira) |
| Syllable repetition | FLRSyl | dan dan dancing (tan tan tanssia) |
| Part-word repetition | FLRSyl | compli- complicated |
| Monosyllabic word repetition | FRLMono | and and and (ja ja ja) |
| Prolongation | FLP | mmmy bag (mmminun laukku) |
| Block | FLB | ...cat (...kissa) |
| Broken word | FLBW | sing...ing (lau...laen) |

**Table 5.**
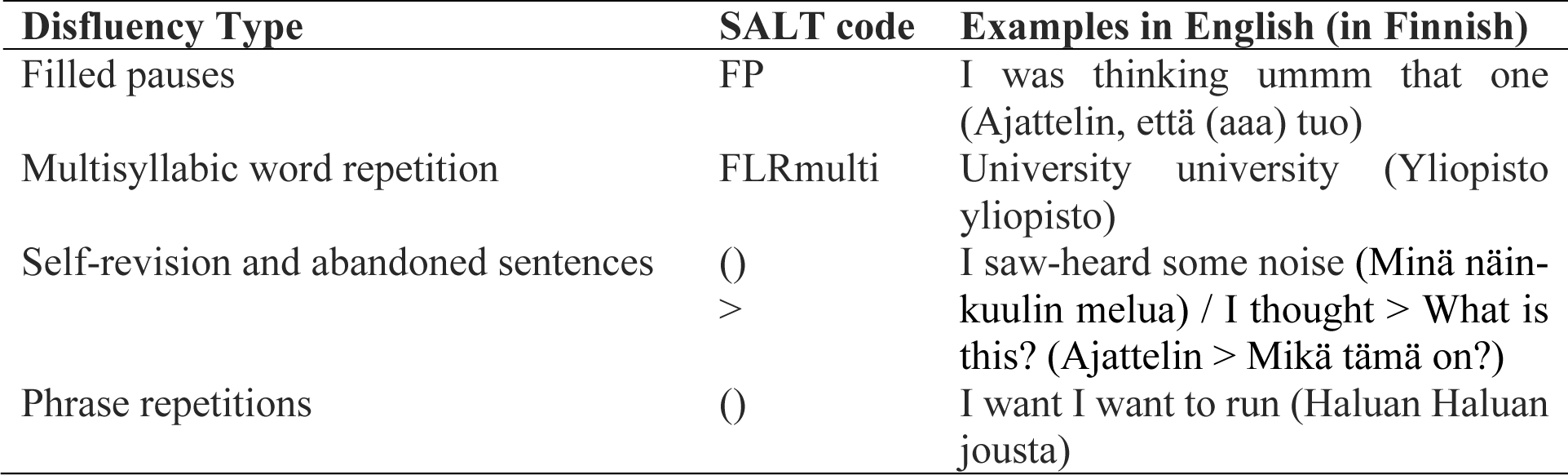
Other types of Disfluencies (ODs)

We calculated disfluency rates using the per 100 words metric. Given the nature of our speech assessments which were focused on conversational speech, we considered a word-based measure more appropriate than the syllable-based metric. Computation of SLDs included combining part word repetition, single-syllable word repetition and dysrhythmic phonation, divided by the word count. For ODs, interjection, revision/abandoned utterances and multi syllable/phrase repetition were combined and then divided by the word count (for a detailed version of these clauses, please refer to Table 1).

The transcriptions were done by a Speech and Language Pathologist specialized in fluency disorders with two master’s level Speech and Language Pathology students. Students were educated in fluency coding and divided the coding of the samples between them. The Fluency Specialized Speech and Language Pathologist was treated as the “golden standard” coder who independently re-coded 10% of each coder’s (speech-language pathologist + two students) transcripts. Point-by-point reliabilities for location were 0.95 (speech language pathologist), 0.95 (student 1) and 0.94 (student 2) between the cross-checked transcripts. Point-by-point reliability for type were 0.90 (speech language pathologist), 0.97 (student 1) and 0.95 (student 2) between the transcripts. For each word in the speech samples, it was determined whether it was spoken fluently or not.

#### Brain Morphology

VBM was the primary derived brain measure of the current study. Additionally, region-of-interest based surface-based analyses of cortical surface area and cortical thickness assessments were conducted to investigate whether these would complement the VBM findings.

Voxel based morphometric analysis of the whole brain was done in SPM12 (Statistical Parametric Mapping) (https://neuro-jena.github.io/cat//index.html#VBM) (Ashburner & Friston, 2000). This procedure was performed to examine differences in brain structure, specifically looking for variations in proportional GMV across the whole brain. The pre-processing was done utilizing the (CAT12) Computational Anatomy Toolbox (http://www.neuro.uni-jena.de/cat/, version r1363) within the SPM framework. A standard preprocessing pipeline was employed, covering normalization, segmentation, quality assessment, and smoothing. The specific details of the pipeline included not splitting the job into separate processes, utilizing the default tissue probability map from SPM, applying affine regularization with the European brains template, choosing medium inhomogeneity correction, using rough affine preprocessing, performing local adaptive segmentation at a medium level, employing skull-stripping through the Graph cuts (GCUT) approach, setting the voxel size for normalized images to 1.5 mm, and fixing internal resampling for preprocessing at 1.0 mm. Dartel was utilized for spatial registration, and both Dartel and Shooting templates were based on the MNI152 template (MNI = Montreal Neurological Institute). Following this, spatial smoothing was performed to enable the application of parametric statistical tests. The images of grey matter underwent smoothing using an 8mm full-width-half-maximum filter (FWHM).

The MRI scans were also preprocessed using the FreeSurfer software (version 6.0). Images with artifacts or large unsegmented regions were removed. These scans were manually edited (Pulli et al., 2022). Once the manual quality control had been performed and manual corrections made, the ENIGMA Cortical Quality Control Protocol 2.0 was employed to conduct a quality check (USC, Mark and Mary Stevens Neuroimaging and Informatics Institute, April 2017), after which some regions were excluded. In total, 34 regions of the brain were parcellated (Supplementary Figure 1).

### Statistical Analyses

#### Whole brain voxel-wise analyses

The SLD and OD distributions were rightly skewed. To address this, we applied a cube root transformation to normalize the distribution of the data (Figure 2). We then conducted a multiple regression analysis using VBM scans to examine the relationship between SLDs and proportional GMV including relevant covariates. The covariates examined in this analysis included the mother’s level of education, the participant’s handedness, age at the time of scanning, and sex of the participant. Maternal education had three levels: Low = Upper secondary school or vocational school or lower, Middle = University of applied sciences, High = University. To simplify the statistical analysis in current study, the low and middle level education of the mother were grouped together, and there were four missing values that were imputed using mode imputation. Handedness, which was measured using parental reports had two levels: right-handed and left-handed, with the four ambidextrous children grouped with right-handed participants to streamline the analysis. Handedness was included as a covariate in the primary model to maintain consistency with prior research (Koenraads et al., 2019). To evaluate the potential impact of reduced power due to low variability in handedness, we conducted an additional analysis excluding this variable (see Supplementary Table 1). Sex was categorized into girls and boys. The interaction effect with sex was also examined. Age was incorporated into the statistical model in terms of days. The same analysis was also done separately for ODs.

**Figure 2.**
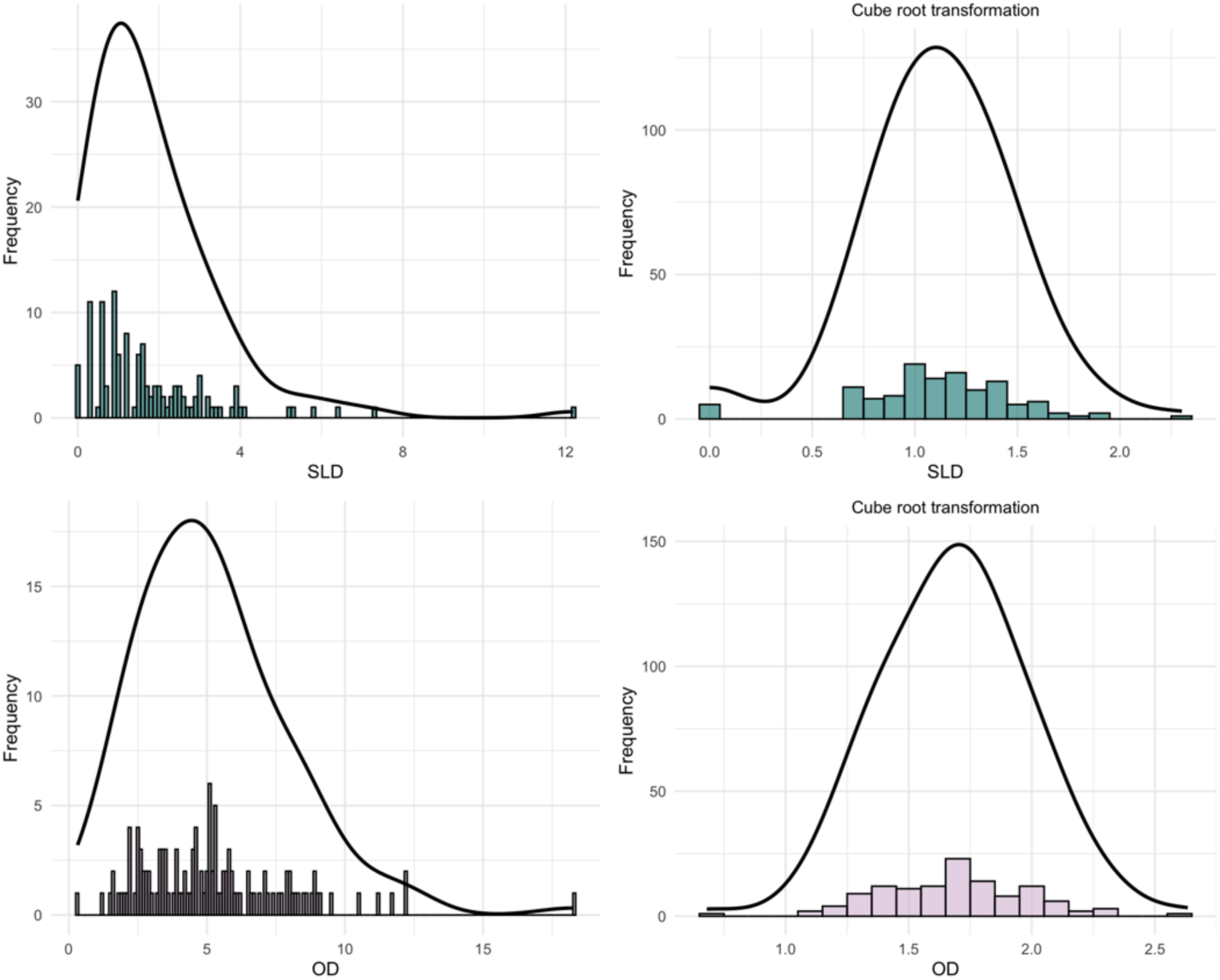
Distribution of speech disfluency measures and their transformations.

The analysis was performed in SPM12 using general linear model (GLM) with “Multiple Regression” on whole-brain data with a threshold of *p* < 0.001 and False Discovery Rate (FDR) corrected cluster level *p*-values < 0.05 were considered statistically significant. The Dartel_AAL atlas (an adaptation of the Automated Anatomical Labeling atlas (Tzourio-Mazoyer et al., 2002) tailored for use with DARTEL-registered images) within SPM was employed to identify areas exhibiting prominent clusters.

#### Complementary region-of-interest based analyses

For surface-based analysis, we extracted measurements of cortical thickness and surface area from the brain’s surface using a tool called aparcstats, which is part of the FreeSurfer software. Desikan-Killiany Atlas (Desikan et al., 2006) was used to identify the regions of the surface-based analysis (Pulli et al., 2022). The statistical analysis was performed in R version 4.4.1 and JASP 0.19.3 using Spearman’s partial correlations between SLD and cortical thickness/surface area in the left MFG and right SFG.

## Results

### Disfluency type and sex differences in disfluency scores

The correlation between SLD and OD in our population was assessed. The results showed a significant positive correlation between the percentage of SLD and OD *r*(118) = 0.53, *p* < .001, 95% CI [0.39, 0.65], indicating a moderate relationship between the two types of disfluencies. This implied that as stuttering-like disfluency increased, other types of disfluencies also tended to increase, suggesting a meaningful relationship between these two measures of disfluency in our sample. Boys had significantly higher SLD values than girls (t (118) = 2.74, *p* = 0.007, 95% CI [0.0517, 0.3204]). Similarly, boys had higher OD values compared to girls (t (118) = 2.58, *p* = 0.011, 95% CI [0.0323, 0.2451]).

### Grey matter volume

A significant positive effect was found between SLDs and proportional GMVs in our 5-year-old participants (Table 4). This positive association was observed in the left Middle Frontal Gyrus (MFG), lobule IX of the left Posterior Cerebellum (PC9) and the right Superior Frontal Gyrus (SFG). No significant interaction effect of sex was observed in the regression model, suggesting that the relationship between SLD and proportional GMVs was consistent across boys and girls. No statistically significant association was found between OD and grey matter.

To assess whether the low variability in handedness affected the results, we compared the primary statistical model (which included handedness as a covariate) with a model excluding it. The resulting clusters remained largely consistent in location, extent, and statistical strength (see Supplementary Table 1), suggesting that the inclusion of handedness did not substantially alter the findings.

**Table 4.**
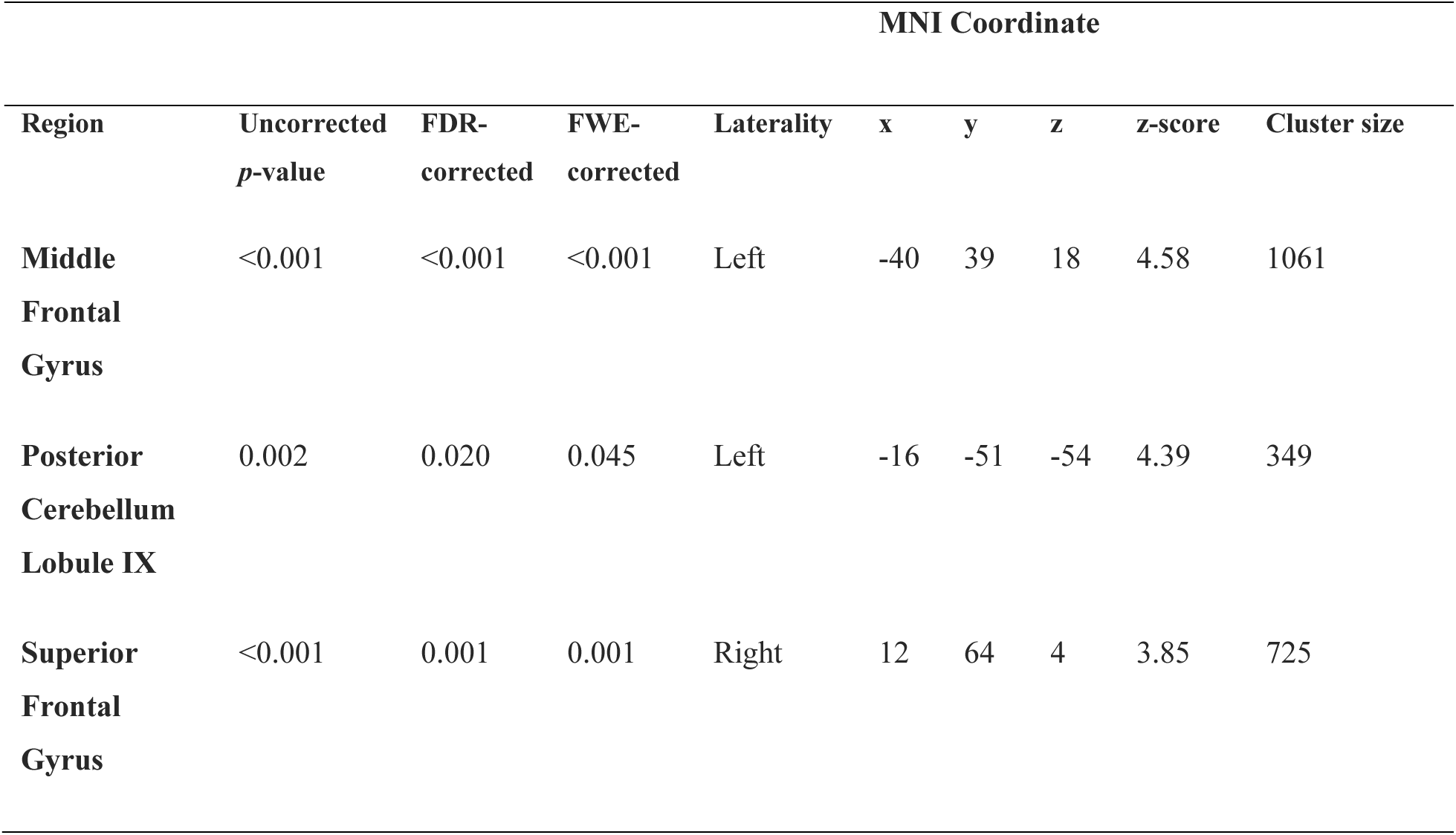
Regions of significant clusters with positive association of SLD and proportional GMVs in children; obtained from the whole-brain VBM analysis.

### Surface-based analysis

A complementary surface-based analysis was conducted using cortical thickness (CT) and surface area (SA) values obtained from FreeSurfer. These metrics were extracted for the regions identified in the VBM analysis, specifically the left MFG and the right SFG. No significant associations were found between the SLD values and either cortical thickness or surface area in these regions (left MFG CT: ρ = −0.037, *p* = .69; left MFG SA: ρ = 0.038, *p* = .68; right SFG CT: ρ = 0.067, *p* = .47; right SFG SA: ρ = 0.055, *p* = .55). Please refer to supplementary material for more information.

## Discussion

We analyzed speech disfluency as a continuum in our non-clinical, typically developing population and investigated whether grey matter characteristics were associated with these measures. Whole-brain analysis revealed significant clusters in the left MFG, PC9 and right SFG, suggesting that these areas are important for speech production. No significant clusters were found in association with OD, indicating that SLD may have distinct neural correlates not shared with other disfluency types. A surface-based analysis examining cortical thickness and surface area as complementary measures to VBM revealed no significant associations with SLD.

### Type of speech disfluency and sex differences

By analyzing SLD and OD as continuous variables across the full sample, rather than categorizing children as stutterers or non-stutterers, we were able to capture the full range of disfluency observed in a typically developing population. This dimensional approach revealed that disfluencies vary gradually rather than occurring exclusively in a clinical subgroup, supporting the view that speech disfluency exists along a continuum (Adams & Runyan, 1981). This aligns with previous research showing that children with stuttering exhibit both stuttering-like and non-stuttering disfluencies in their speech (Ambrose & Yairi, 1999; Johnson, 1959; Tumanova et al., 2014; Yairi & Ambrose, 2005). The positive relationship observed between SLD and OD in our sample further supports this notion, suggesting that these behaviors co-occur along a shared continuum of fluency – yet they might have differential neural correlates.

Consistent with broader theories emphasizing the role of motor and cognitive control processes in fluent speech production, we found that SLD was associated with structural differences in the left MFG and PC9. Although not classically highlighted in early speech motor control models e.g., (Guenther, 2016), the left anterior MFG has been implicated in speech monitoring, planning, and executive aspects of verbal fluency tasks (Abrahams et al., 2003; Turken & Dronkers, 2011). Similarly, the posterior cerebellum has been increasingly recognized for its role in motor timing and coordination relevant to speech fluency (Ackermann, 2008). The involvement of motor-control circuits in our findings aligns with broader lesion-mapping evidence (Theys et al., 2024) implicating subcortical motor systems in clinical stuttering, suggesting that even subclinical disfluencies engage key neural circuits underlying speech motor control. In contrast, the anticipated association between OD and language-related cortex (inferior frontal, parietal, temporal regions) was not supported, we found no significant structural links for OD, failing to confirm the hypothesized language-network involvement from Hickok and Poeppel’s (2007) dual-stream model. Although SLD and OD were moderately correlated, the neural associations appeared more robust for SLD, suggesting that different disfluency types may engage overlapping but partially distinct brain mechanisms. By modeling SLD and OD as separate dimensions, we were able to highlight the value of treating disfluency subtypes separately rather than as a single, homogeneous construct.

Our findings also revealed that boys exhibited significantly more disfluencies (both SLD and OD) than girls. This finding is consistent with previous research highlighting a higher prevalence of stuttering in boys compared to girls (Proctor et al., 2008). While the current study focuses on speech disfluencies rather than clinically diagnosed stuttering, these results align with broader patterns observed in speech and language development, where boys are often found to experience more challenges than girls (Anastasi, 1937; *The Development of Sex Differences*, 1966; Maccoby & Jacklin, 1978). However, it is important to note that the magnitude of sex differences in speech and language development remains debated. While some studies report significant disparities favoring boys (Proctor et al., 2008), others suggest these differences are modest or negligible (Hyde, 2005, 2016; Lindberg et al., 2010; Zell et al., 2015). For instance, the developmental trajectory of speech and language abilities may differ across ages, making it challenging to draw definitive conclusions about the long-term significance of sex differences at any single developmental stage (Etchell et al., 2018).

### Grey matter association with speech disfluency

We found a large cluster in the left anterior MFG, which approximately maps to Brodmann area 46 (BA46) of the dorsolateral prefrontal cortex (DLPFC) (Figure 3B). This falls anatomically outside the classical “Broca’s area,” which is confined to the inferior frontal gyrus (BA44 and BA45) (Binkofski & Buccino, 2004). BA46 of the anterior MFG is found to subserve domain-general executive operations like working memory, planning, and cognitive control. Though not a core perisylvian language area, the left anterior MFG is often active during language tasks with higher-order processing (Hertrich et al., 2021). For example, this DLPFC region supports word retrieval in verbal fluency tasks (Abrahams et al., 2003) and the phonological storage component of working memory in speech tasks (Wen et al., 2017). It has also been involved in the broader language network for both language production (e.g. meta-analytic connectivity studies show BA46 is engaged in the speech production system (Ardila et al., 2016)) and language comprehension (lesion mapping studies in aphasia show BA46 is one of the regions critical for comprehension of language (Turken & Dronkers, 2011)). Earlier research on stuttering and disfluency has also involved the left MFG/DLPFC in the regulation of fluency. (Chang et al., 2008) conducted a VBM study comparing children with persistent stuttering, those recovered from stuttering, and fluent controls. Their findings revealed reduced grey matter volume in the left MFG in recovered stutterers compared to persistent stutterers. This suggests that structural anomalies in the left MFG may serve as a neuroanatomical marker associated with speech disfluency.

**Figure 3.**
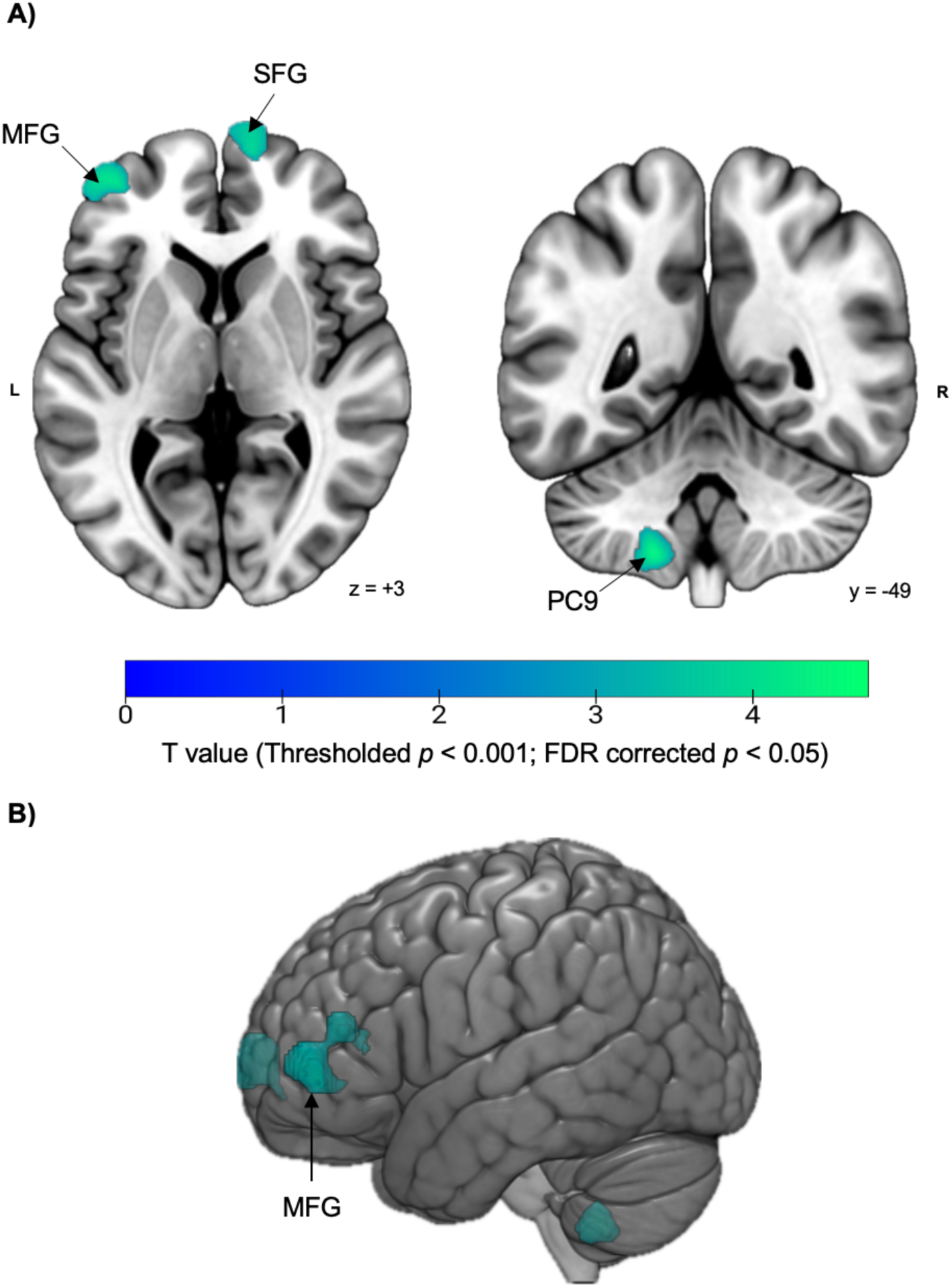
**A)** Clusters signify the positive association of SLD with proportional GMV. The color bar represents the T-scores of the SLD cluster values in the entire population. MFG = Middle Frontal Gyrus; SFG = Superior Frontal Gyrus; PC9 = Posterior Cerebellum Lobule IX. **B)** Rendered 3D view of the significant cluster in the left anterior middle frontal gyrus (MFG) associated with higher levels of SLD. The cluster is displayed on the lateral surface of the left hemisphere. This region corresponds approximately to Brodmann area 46.

We also identified a significant cluster in the left posterior cerebellum, specifically in lobule IX. The cerebellum, traditionally recognized for its role in motor coordination, has increasingly been implicated in cognitive functions, including language and speech (E et al., 2014; Hogan et al., 2011; Stoodley & Schmahmann, 2009). Research indicates that the posterior cerebellum, particularly lobules VI–IX, is engaged in cognitive processes such as working memory, language, and reading in both adults and children (Grogan et al., 2009; He et al., 2013; Kronbichler et al., 2008; Richardson & Price, 2009). Moreover, studies have demonstrated that increased grey matter in these regions is associated with better cognitive performance in children, including vocabulary, reading, and executive functioning (Moore et al., 2017).

Building on the involvement of the left MFG and PC9 in speech disfluency, our analysis also revealed a significant cluster in the right SFG. This finding extends previous research, which has largely focused on left-hemisphere contributions to speech and language processing and highlights the importance of right hemisphere areas in speech disfluency. The right SFG is a region not frequently highlighted in studies of speech and language processing. Most prior research has focused on the left SFG and its connections, particularly in the context of speech initiation and language recovery. For example, a study demonstrated a corticocortical network between the left SFG and Broca’s area, suggesting a role in motor and language integration (Ookawa et al., 2017). Similarly, another study found that theta-burst stimulation targeting the left SFG significantly improved language recovery in post-stroke aphasia patients, underscoring its potential as a language-related hub (Ren et al., 2023).

While these studies provide valuable insights, they primarily focus on functional or structural connectivity, rather than proportional GMV, and predominantly examine the left hemisphere. Our finding in the right SFG extends this body of work by implicating the contralateral region; however, given that right SFG has not been consistently implicated in stuttering-related research, caution is warranted in interpreting its role. It is possible that the right SFG cluster reflects broader cognitive processes, such as attention and executive control, which may indirectly support fluent speech rather than being specific to disfluency per se. This interpretation is consistent with the notion that the right hemisphere, while less directly associated with core language production, contributes to cognitive and motor integration necessary for speech (Bernard et al., 2018).

### Lack of associations in surface-based metrics

While our VBM analysis identified significant GMV differences related to SLD, we did not observe corresponding changes in cortical thickness or surface area. This may reflect the subtler nature of neural variation in our non-clinical, population-based sample. Unlike studies which examined children with clinically diagnosed persistent stuttering and reported surface-based differences in motor and frontal regions (Garnett et al., 2018; Koenraads et al., 2019), our participants exhibited naturally occurring variation in disfluency without a clinical label.

It is possible that the structural differences observed in our study reflect functional or microarchitectural adaptations that do not produce gross anatomical changes in cortical thickness or surface area. VBM is sensitive to fine-grained variations in tissue composition and local density, which may not always manifest in surface-based measures (Goto et al., 2022). These findings highlight the value of multimodal approaches and underscore the importance of considering ongoing brain maturation and structural variability during early childhood when interpreting associations with speech disfluency.

### Strengths and limitations

One of the major strengths of the current study is that we treated the disfluency variable as a continuous measure instead of dividing the participants into stuttering and non-stuttering groups. This approach allowed us to investigate morphological brain differences in individuals with different types of speech disfluencies, irrespective of whether they fit into a specific diagnostic category. We conducted our research on 5-year-old participants, which can be considered as an appropriate age to identify structural brain anomalies for speech disruptions. We also had a sizeable sample of 120 children with usable speech and neuroimaging data, which could aid future research in exploring the structural similarities and differences in the brains of individuals with speech difficulties. Overall, our study has the potential to contribute to the literature on speech disfluencies. Along with that, this study has several limitations that should be acknowledged. First, the cross-sectional design limits our ability to infer causal relationships between structural brain differences and speech disfluency. Longitudinal studies are necessary to explore how these relationships develop over time. Second, the findings rely on specific atlases for computing CT and SA (Desikan-Killiany atlas), which may limit comparability with other studies using different parcellation schemes. Finally, the classification strategy of speech disfluency could influence the findings and their interpretation.

### Conclusion

This study identifies key neural correlates of speech disfluency in children, including the left middle frontal gyrus, left posterior cerebellum, and right superior frontal gyrus. These findings underscore the involvement of both hemispheres and the integration of motor and cognitive neural networks in speech fluency. Notably, our dimensional approach – without categorizing children as stutterers or non-stutterers – identified structural differences in brain regions that have also been implicated in clinical studies of stuttering. This convergence suggests that natural variation in speech disfluency engages similar neural systems, supporting the relevance of a continuum-based perspective in both research and clinical contexts.

## Acknowledgements

We are grateful to the participants of the FinnBrain Birth Cohort study, as well as the staff and support personnel involved. Special thanks to research nurse Susanne Sinisalo, neuroradiologist Riitta Parkkola, paediatric neurologist Tuire Lähdesmäki, and physicist Jani Saunavaara for their contributions to the 5-year MRI data collection. We also thank the University of Turku Speech-Language Pathology students for their assistance in data collection and coding as well as SLP Päivi Kantomaa for conducting control transcriptions, which helped us ensure the reliability of the speech sample transcriptions.

## Author Contributions

- AJ: data and statistical analysis, leading the manuscript writing
- AYS and EsS: speech data collection, coding of speech samples
- EPP: MRI data collection, VBM and FreeSurfer pre-processing
- HR: drafting initial version of manuscript
- HM, AC, VK, EkS and EeS: MRI data collection
- LK: established the FinnBrain Birth Cohort and funded FinnBrain cohort infrastructure
- HK: established the FinnBrain Birth Cohort and funded the MRI scans.
- NJ: funding acquisition, supervision
- EMA: conceptualization of the study, drafting initial version of manuscript, supervision
- JJT: conceptualization of this study, funding acquisition, leading the data collection, implementation and supervision of MRI data analysis, drafting initial version of manuscript, supervision of AJ

## All authors contributed to critical revision of the manuscript. All authors read and approved the final manuscript

## Data availability statement

The Finnish law and ethical permissions do not allow open sharing of the data used in this study, but data access is possible via formal material transfer agreements (MTA). Investigators that wish to access the data are encouraged to contact Principal Investigator of the FinnBrain Birth Cohort study Linnea Karlsson.

## References

Abrahams, S., Goldstein, L. H., Simmons, A., Brammer, M. J., Williams, S. C. R., Giampietro, V. P., Andrew, C. M., & Leigh, P. N. (2003). Functional magnetic resonance imaging of verbal fluency and confrontation naming using compressed image acquisition to permit overt responses. Human Brain Mapping, 20(1), 29–40. 10.1002/hbm.10126

Ackermann, H. (2008). Cerebellar contributions to speech production and speech perception: Psycholinguistic and neurobiological perspectives. Trends in Neurosciences, 31(6), 265–272. 10.1016/j.tins.2008.02.011

Adams, M. R., & Runyan, C. M. (1981). Stuttering and fluency: Exclusive events or points on a continuum? Journal of Fluency Disorders, 6(3), 197–218. 10.1016/0094-730X(81)90002-4

Ambrose, N. G., & Yairi, E. (1999). Normative disfluency data for early childhood stuttering. *Journal of Speech*, Language, and Hearing Research: JSLHR, 42(4), 895–909. 10.1044/jslhr.4204.895

Anastasi, A. (1937). Differential psychology. Macmillan.

Ardila, A., Bernal, B., & Rosselli, M. (2016). Connectivity of BA46 involvement in the executive control of language. Psicothema, 28(1), 26–31.

ASHA. (2024). Fluency Disorders. American Speech-Language-Hearing Association; American Speech-Language-Hearing Association. https://www.asha.org/practice-portal/clinical-topics/fluency-disorders/

Ashburner, J., & Friston, K. J. (2000). Voxel-based morphometry—The methods. NeuroImage, 11(6 Pt 1), 805–821. 10.1006/nimg.2000.0582

Beal, D. S., Gracco, V. L., Brettschneider, J., Kroll, R. M., & De Nil, L. F. (2013). A voxel-based morphometry (VBM) analysis of regional grey and white matter volume abnormalities within the speech production network of children who stutter. Cortex; a Journal Devoted to the Study of the Nervous System and Behavior, 49(8), 2151–2161. 10.1016/j.cortex.2012.08.013

Beal, D. S., Gracco, V. L., Lafaille, S. J., & De Nil, L. F. (2007). Voxel-based morphometry of auditory and speech-related cortex in stutterers. Neuroreport, 18(12), 1257–1260. 10.1097/WNR.0b013e3282202c4d

Bernard, F., Lemée, J.-M., Ter Minassian, A., & Menei, P. (2018). Right Hemisphere Cognitive Functions: From Clinical and Anatomic Bases to Brain Mapping During Awake Craniotomy Part I: Clinical and Functional Anatomy. World Neurosurgery, 118, 348–359. 10.1016/j.wneu.2018.05.024

Binkofski, F., & Buccino, G. (2004). Motor functions of the Broca’s region. Brain and Language, 89(2), 362–369. 10.1016/S0093-934X(03)00358-4

Boberg, E., & Kully, D. (1994). Long-term results of an intensive treatment program for adults and adolescents who stutter. Journal of Speech and Hearing Research, 37(5), 1050–1059. 10.1044/jshr.3705.1050

Boey, R. A., Wuyts, F. L., Van de Heyning, P. H., De Bodt, M. S., & Heylen, L. (2007). Characteristics of stuttering-like disfluencies in Dutch-speaking children. Journal of Fluency Disorders, 32(4), 310–329. 10.1016/j.jfludis.2007.07.003

Carlo, E. J., & Watson, J. B. (2003). Disfluencies of 3- and 5-year old Spanish-speaking children. Journal of Fluency Disorders, 28(1), 37–53. 10.1016/S0094-730X(03)00004-4

Chang, S.-E., Erickson, K. I., Ambrose, N. G., Hasegawa-Johnson, M. A., & Ludlow, C. L. (2008). Brain anatomy differences in childhood stuttering. NeuroImage, 39(3), 1333– 1344. 10.1016/j.neuroimage.2007.09.067

Chow, H. M., Garnett, E. O., Koenraads, S. P. C., & Chang, S.-E. (2023). Brain developmental trajectories associated with childhood stuttering persistence and recovery. Developmental Cognitive Neuroscience, 60, 101224. 10.1016/j.dcn.2023.101224

Conture, E. G. (2001). Stuttering: Its Nature, Diagnosis, and Treatment. Allyn and Bacon.

Desikan, R. S., Ségonne, F., Fischl, B., Quinn, B. T., Dickerson, B. C., Blacker, D., Buckner, R. L., Dale, A. M., Maguire, R. P., Hyman, B. T., Albert, M. S., & Killiany, R. J. (2006). An automated labeling system for subdividing the human cerebral cortex on MRI scans into gyral based regions of interest. NeuroImage, 31(3), 968–980. 10.1016/j.neuroimage.2006.01.021

E, K.-H., Chen, S.-H. A., Ho, M.-H. R., & Desmond, J. E. (2014). A meta-analysis of cerebellar contributions to higher cognition from PET and fMRI studies. Human Brain Mapping, 35(2), 593–615. 10.1002/hbm.22194

Etchell, A., Adhikari, A., Weinberg, L. S., Choo, A. L., Garnett, E. O., Chow, H. M., & Chang, S.-E. (2018). A Systematic Literature Review of Sex Differences inChildhood Language and Brain Development. Neuropsychologia, 114, 19–31. 10.1016/j.neuropsychologia.2018.04.011

Fischl, B. (2012). FreeSurfer. NeuroImage, 62(2), 774–781. 10.1016/j.neuroimage.2012.01.021

Garnett, E. O., Chow, H. M., Nieto-Castañón, A., Tourville, J. A., Guenther, F. H., & Chang, S.-E. (2018). Anomalous morphology in left hemisphere motor and premotor cortex of children who stutter. Brain: A Journal of Neurology, 141(9), 2670–2684. 10.1093/brain/awy199

Goto, M., Abe, O., Hagiwara, A., Fujita, S., Kamagata, K., Hori, M., Aoki, S., Osada, T., Konishi, S., Masutani, Y., Sakamoto, H., Sakano, Y., Kyogoku, S., & Daida, H. (2022). Advantages of Using Both Voxel- and Surface-based Morphometry in Cortical Morphology Analysis: A Review of Various Applications. Magnetic Resonance in Medical Sciences, 21(1), 41–57. 10.2463/mrms.rev.2021-0096

Grogan, A., Green, D. W., Ali, N., Crinion, J. T., & Price, C. J. (2009). Structural Correlates of Semantic and Phonemic Fluency Ability in First and Second Languages. Cerebral Cortex, 19(11), 2690–2698. 10.1093/cercor/bhp023

Guenther, F. H. (2016). Neural Control of Speech. The MIT Press. 10.7551/mitpress/10471.001.0001

He, Q., Xue, G., Chen, C., Chen, C., Lu, Z.-L., & Dong, Q. (2013). Decoding the Neuroanatomical Basis of Reading Ability: A Multivoxel Morphometric Study. Journal of Neuroscience, 33(31), 12835–12843. 10.1523/JNEUROSCI.0449-13.2013

Hertrich, I., Dietrich, S., Blum, C., & Ackermann, H. (2021). The Role of the Dorsolateral Prefrontal Cortex for Speech and Language Processing. Frontiers in Human Neuroscience, 15, 645209. 10.3389/fnhum.2021.645209

Hickok, G., & Poeppel, D. (2007). The cortical organization of speech processing. Nature Reviews Neuroscience, 8(5), 393–402. 10.1038/nrn2113

Hogan, M. J., Staff, R. T., Bunting, B. P., Murray, A. D., Ahearn, T. S., Deary, I. J., & Whalley, L. J. (2011). Cerebellar brain volume accounts for variance in cognitive performance in older adults. Cortex; a Journal Devoted to the Study of the Nervous System and Behavior, 47(4), 441–450. 10.1016/j.cortex.2010.01.001

Hyde, J. S. (2005). The gender similarities hypothesis. American Psychologist, 60(6), 581–592. 10.1037/0003-066X.60.6.581

Hyde, J. S. (2016). Sex and cognition: Gender and cognitive functions. Current Opinion in Neurobiology, 38, 53–56. 10.1016/j.conb.2016.02.007

Jäncke, L., Hänggi, J., & Steinmetz, H. (2004). Morphological brain differences between adult stutterers and non-stutterers. BMC Neurology, 4(1), 23. 10.1186/1471-2377-4-23

Jansson-Verkasalo, E., Silvén, M., Lehtiö, I., & Eggers, K. (2021). Speech disfluencies in typically developing Finnish-speaking children – preliminary results. Clinical Linguistics & Phonetics, 35(8), 707–726. 10.1080/02699206.2020.1818287

Johnson, W. (1959). The onset of stuttering: Research findings and implications. University of Minnesota Press.

Karlsson, L., Tolvanen, M., Scheinin, N. M., Uusitupa, H.-M., Korja, R., Ekholm, E., Tuulari, J. J., Pajulo, M., Huotilainen, M., Paunio, T., Karlsson, H., & Group, F. B. C. S. (2018). Cohort Profile: The FinnBrain Birth Cohort Study (FinnBrain). International Journal of Epidemiology, 47(1), 15–16j. 10.1093/ije/dyx173

Kell, C. A., Neumann, K., von Kriegstein, K., Posenenske, C., von Gudenberg, A. W., Euler, H., & Giraud, A.-L. (2009). How the brain repairs stuttering. Brain, 132(10), 2747–2760. 10.1093/brain/awp185

Koenraads, S. P. C., El Marroun, H., Muetzel, R. L., Chang, S. E., Vernooij, M. W., Baatenburg de Jong, R. J., White, T., Franken, M. C., & van der Schroeff, M. P. (2019). Stuttering and gray matter morphometry: A population-based neuroimaging study in young children. Brain and Language, 194, 121–131. 10.1016/j.bandl.2019.04.008

Kronbichler, M., Wimmer, H., Staffen, W., Hutzler, F., Mair, A., & Ladurner, G. (2008). Developmental dyslexia: Gray matter abnormalities in the occipitotemporal cortex. Human Brain Mapping, 29(5), 613–625. 10.1002/hbm.20425

Lindberg, S. M., Hyde, J. S., Petersen, J. L., & Linn, M. C. (2010). New trends in gender and mathematics performance: A meta-analysis. Psychological Bulletin, 136(6), 1123–1135. 10.1037/a0021276

Lu, C., Peng, D., Chen, C., Ning, N., Ding, G., Li, K., Yang, Y., & Lin, C. (2010). Altered effective connectivity and anomalous anatomy in the basal ganglia-thalamocortical circuit of stuttering speakers. Cortex; a Journal Devoted to the Study of the Nervous System and Behavior, 46(1), 49–67. 10.1016/j.cortex.2009.02.017

Maccoby, E. E. (Ed.). (1966). The development of sex differences. Stanford University Press.

Maccoby, E. E., & Jacklin, C. N. (1978). The Psychology of Sex Differences: —Vol. II: Annotated Bibliography. Stanford University Press.

Miller, J., & Iglesias, A. (2020). Systematic Analysis of Language Transcripts (SALT) (Version 20) [Computer software].

Moore, D. M., D’Mello, A. M., McGrath, L. M., & Stoodley, C. J. (2017). The developmental relationship between specific cognitive domains and grey matter in the cerebellum. Developmental Cognitive Neuroscience, 24, 1–11. 10.1016/j.dcn.2016.12.001

Natke, U., Sandrieser, P., Pietrowsky, R., & Kalveram, K. T. (2006). Disfluency data of German preschool children who stutter and comparison children. Journal of Fluency Disorders, 31(3), 165–176. 10.1016/j.jfludis.2006.04.002

Ookawa, S., Enatsu, R., Kanno, A., Ochi, S., Akiyama, Y., Kobayashi, T., Yamao, Y., Kikuchi, T., Matsumoto, R., Kunieda, T., & Mikuni, N. (2017). Frontal Fibers Connecting the Superior Frontal Gyrus to Broca Area: A Corticocortical Evoked Potential Study. World Neurosurgery, 107, 239–248. 10.1016/j.wneu.2017.07.166

Proctor, A., Yairi, E., Duff, M. C., & Zhang, J. (2008). Prevalence of stuttering in African American preschoolers. Journal of Speech, Language, and Hearing Research: JSLHR, 51(6), 1465–1479. 10.1044/1092-4388(2008/07-0057)

Pulli, E. P., Silver, E., Kumpulainen, V., Copeland, A., Merisaari, H., Saunavaara, J., Parkkola, R., Lähdesmäki, T., Saukko, E., Nolvi, S., Kataja, E.-L., Korja, R., Karlsson, L., Karlsson, H., & Tuulari, J. J. (2022). Feasibility of FreeSurfer Processing for T1-Weighted Brain Images of 5-Year-Olds: Semiautomated Protocol of FinnBrain Neuroimaging Lab. Frontiers in Neuroscience, 16. 10.3389/fnins.2022.874062

Ren, J., Ren, W., Zhou, Y., Dahmani, L., Duan, X., Fu, X., Wang, Y., Pan, R., Zhao, J., Zhang, P., Wang, B., Yu, W., Chen, Z., Zhang, X., Sun, J., Ding, M., Huang, J., Xu, L., Li, S., … Liu, H. (2023). Personalized functional imaging-guided rTMS on the superior frontal gyrus for post-stroke aphasia: A randomized sham-controlled trial. Brain Stimulation, 16(5), 1313–1321. 10.1016/j.brs.2023.08.023

Richardson, F. M., & Price, C. J. (2009). Structural MRI studies of language function in the undamaged brain. Brain Structure and Function, 213(6), 511–523. 10.1007/s00429-009-0211-y

Schmidt, R. (1992). Psychological Mechanisms Underlying Second Language Fluency. Studies in Second Language Acquisition, 14(4), 357–385. 10.1017/S0272263100011189

Smith, A., & Weber, C. (2017). How Stuttering Develops: The Multifactorial Dynamic Pathways Theory. *Journal of Speech*, Language, and Hearing Research, 60(9), 2483–2505. 10.1044/2017_JSLHR-S-16-0343

Sowman, P. F., Ryan, M., Johnson, B. W., Savage, G., Crain, S., Harrison, E., Martin, E., & Burianová, H. (2017). Grey matter volume differences in the left caudate nucleus of people who stutter. Brain and Language, 164, 9–15. 10.1016/j.bandl.2016.08.009

Stoodley, C. J., & Schmahmann, J. D. (2009). Functional topography in the human cerebellum: A meta-analysis of neuroimaging studies. NeuroImage, 44(2), 489–501. 10.1016/j.neuroimage.2008.08.039

Theys, C., Jaakkola, E., Melzer, T. R., De Nil, L. F., Guenther, F. H., Cohen, A. L., Fox, M. D., & Joutsa, J. (2024). Localization of stuttering based on causal brain lesions. Brain: A Journal of Neurology, 147(6), 2203–2213. 10.1093/brain/awae059

Tichenor, S. E., & Yaruss, J. S. (2021). Variability of Stuttering: Behavior and Impact. American Journal of Speech-Language Pathology, 30(1), 75–88. 10.1044/2020_AJSLP-20-00112

Tumanova, V., Conture, E. G., Lambert, E. W., & Walden, T. A. (2014). Speech disfluencies of preschool-age children who do and do not stutter. Journal of Communication Disorders, 49, 25–41. 10.1016/j.jcomdis.2014.01.003

Turken, A. U., & Dronkers, N. F. (2011). The Neural Architecture of the Language Comprehension Network: Converging Evidence from Lesion and Connectivity Analyses. Frontiers in Systems Neuroscience, 5, 1. 10.3389/fnsys.2011.00001

Wen, J., Yu, Tao, Liu, Li, Hu, Zhenhong, Yan, Jiaqing, Li, Yongjie, & and Li, X. (2017). Evaluating the roles of left middle frontal gyrus in word production using electrocorticography. Neurocase, 23(5–6), 263–269. 10.1080/13554794.2017.1387275

Yairi, E., & Ambrose, N. G. (2005). Early Childhood Stuttering for Clinicians by Clinicians. PRO-ED.

Zell, E., Krizan, Z., & Teeter, S. R. (2015). Evaluating gender similarities and differences using metasynthesis. American Psychologist, 70(1), 10–20. 10.1037/a0038208

