## Supplementary_Material for "Structural Brain Correlates of Speech Disfluency in Early Childhood: A Dimensional Analysis in a Non-Clinical Cohort"

Statistical model results without including the handedness variable:

**Supplementary Table 1**. Regions of significant clusters with positive association of SLD and grey matter densities in children; obtained from the whole-brain voxel-based morphometry analysis (without the handedness variable).

|  |  |  |  |  | **MNI Coordinate** | | |  |  |
| --- | --- | --- | --- | --- | --- | --- | --- | --- | --- |
| **Region** | **Uncorrected *p*-value** | **FDR-corrected** | **FWE-corrected** | **Laterality** | **x** | **y** | **z** | **z-score** | **Cluster size** |
| **Middle Frontal Gyrus** | <0.001 | <0.001 | <0.001 | Left | -40 | 39 | 18 | 4.61 | 1113 |
| **Posterior Cerebellum Lobule IX** | 0.002 | 0.019 | 0.043 | Left | -16 | -51 | -54 | 4.40 | 353 |
| **Superior Frontal Gyrus** | <0.001 | 0.001 | 0.001 | Right | 12 | 64 | 4 | 3.83 | 716 |

*FreeSurfer Processing*

FreeSurfer version 6.0.0 (https://surfer.nmr.mgh.harvard.edu/) was used to process T1-weighted brain images of 5-year-old children, performing semiautomated segmentation of cortical metrics such as cortical thickness and surface area. Cortical metrics were extracted using the "aparcstats" function, which computes region-wise statistics based on the Desikan-Killiany atlas. The study employed additional quality control procedures, including manual edits and a dichotomous quality rating system, to address segmentation errors caused by motion artifacts. While manual edits minimally affected cortical thickness results (less than 2%), quality checks were critical for reliable data. For more information, please see (Pulli et al., 2022).

(A)
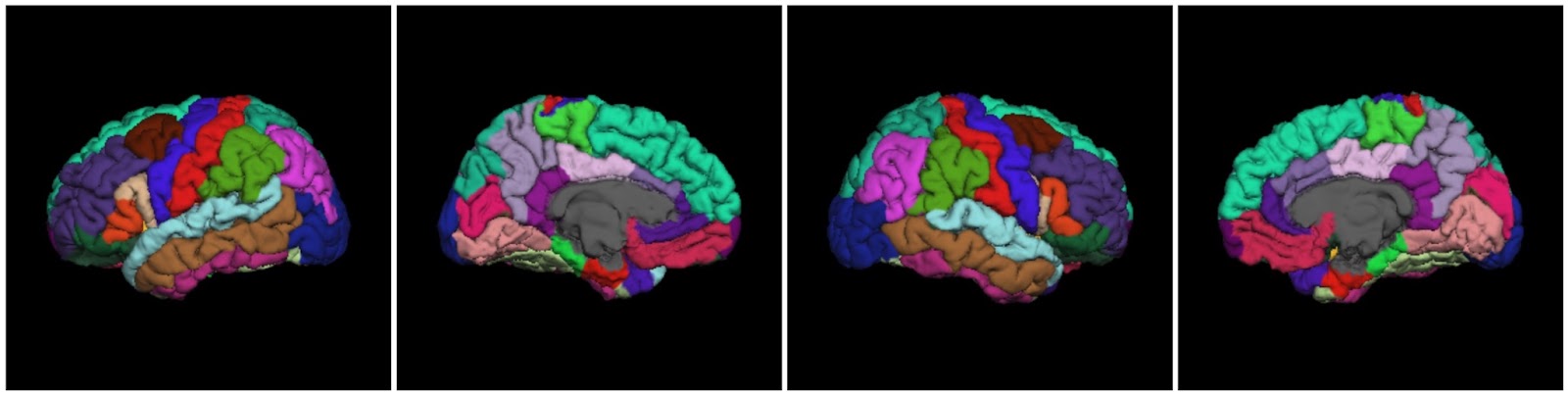
(B)

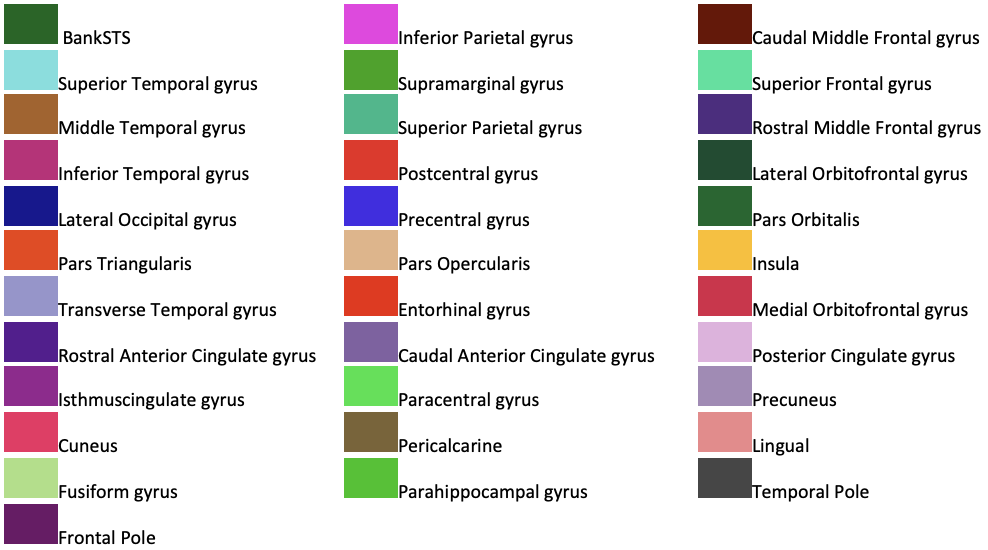

**Supplementary Figure 1**. (A) Using the Desikan-Killiany Atlas and the ENIGMA quality control protocol, the 34 parcellated regions are identified and color coded for one of our 5-year-old participants; Image Source: USC, Mark and Mary Stevens Neuroimaging and Informatics Institute, April 2017; (B) Color-coded references for the regions

*Statistical analysis*

R version 4.4.1 and JASP 0.19.3 were used to perform the statistical analysis. We focused on two regions of interest to explore whether they could complement the VBM findings: the left middle frontal gyrus (MFG) and the right superior frontal gyrus (SFG). Among the parcellated regions, we specifically selected the right SFG and left rostral MFG, as these corresponded to areas where significant clusters were observed. Cortical thickness and surface area measures were extracted for these regions, although some data were missing for certain participants in the study.

Missing data for cortical thickness and surface area measures were addressed using two different imputation methods based on the extent of missingness. For the left rostral MFG, where only four values were missing, mean imputation was performed. This approach was deemed appropriate given the minimal proportion of missing values, ensuring the dataset's integrity without introducing significant bias. For the right SFG, with 31 missing values, multiple imputation using chained equations (MICE) was implemented (Buuren & Groothuis-Oudshoorn, 2011). The MICE procedure employed predictive mean matching (PMM) as the imputation method, incorporating age at scan and sex as predictors to improve estimation accuracy. Five imputed datasets were generated with a maximum of 50 iterations, and the first imputed dataset was selected for analysis. This method provided robust imputed values by leveraging the relationships between cortical thickness/surface area and the chosen predictors while accommodating the higher proportion of missing data.

We replicated the VBM analysis by including the same covariates used in the SPM model. We conducted Spearman’s partial correlation (as these values were not normally distributed) for both cortical thickness and surface area of the left MFG and the right SFG.

**Supplementary Table 2.**

| *Spearman's Partial Correlations: Cortical thickness of left MFG and SLD* | | | | | | |
| --- | --- | --- | --- | --- | --- | --- |
| Variable | |  | | SLD | | Left MFG |
| 1. SLD |  | n |  | — |  |  |
|  |  | Spearman's rho |  | — |  |  |
|  |  | p-value |  | — |  |  |
| 2. Left MFG |  | n |  | 120 |  | — |
|  |  | Spearman's rho |  | -0.037 |  | — |
|  |  | p-value |  | 0.696 |  | — |
| *Note.* Conditioned on variables: sex, mother’s education, age, handedness. | | | | | | |
| * p < .05, ** p < .01, *** p < .001 | | | | | | |

**Supplementary Table 3.**

| *Spearman's Partial Correlations: Cortical thickness of right SFG and SLD* | | | | | | |
| --- | --- | --- | --- | --- | --- | --- |
| Variable | |  | | SLD | | Right SFG |
| 1. SLD |  | n |  | — |  |  |
|  |  | Spearman's rho |  | — |  |  |
|  |  | p-value |  | — |  |  |
| 2. Right SFG |  | n |  | 120 |  | — |
|  |  | Spearman's rho |  | 0.067 |  | — |
|  |  | p-value |  | 0.477 |  | — |
| *Note.* Conditioned on variables: sex, mother’s education, age, handedness. | | | | | | |
| * p < .05, ** p < .01, *** p < .001 | | | | | | |

**Supplementary Table 4.**

| *Spearman's Partial Correlations: Surface Area of Left MFG with SLD* | | | | | | |
| --- | --- | --- | --- | --- | --- | --- |
| Variable | |  | | SLD | | Left MFG |
| 1. SLD |  | n |  | — |  |  |
|  |  | Spearman's rho |  | — |  |  |
|  |  | p-value |  | — |  |  |
| 2. Left MFG |  | n |  | 120 |  | — |
|  |  | Spearman's rho |  | 0.038 |  | — |
|  |  | p-value |  | 0.685 |  | — |
| *Note.* Conditioned on variables: sex, mother’s education, age, handedness. | | | | | | |
| * p < .05, ** p < .01, *** p < .001 | | | | | | |

**Supplementary Table 5.**

| *Spearman's Partial Correlations: Surface Area of Right SFG with SLD* | | | | | | |
| --- | --- | --- | --- | --- | --- | --- |
| Variable | |  | | SLD | | Right SFG |
| 1. SLD |  | n |  | — |  |  |
|  |  | Spearman's rho |  | — |  |  |
|  |  | p-value |  | — |  |  |
| 2. Right SFG |  | n |  | 120 |  | — |
|  |  | Spearman's rho |  | 0.055 |  | — |
|  |  | p-value |  | 0.556 |  | — |
| *Note.* Conditioned on variables: sex, mother’s education, age, handedness. | | | | | | |
| * p < .05, ** p < .01, *** p < .001 | | | | | | |

As shown in the results, there is no significant relationship between SLD and the cortical thickness or the surface area of either the left MFG or the right SFG. This indicates that, once accounting for other variables, there is no statistically meaningful association between these two factors and SLD for these areas.

**References**

Buuren, S. van, & Groothuis-Oudshoorn, K. (2011). mice: Multivariate Imputation by Chained Equations in R. *Journal of Statistical Software*, *45*, 1–67. https://doi.org/10.18637/jss.v045.i03

Pulli, E. P., Silver, E., Kumpulainen, V., Copeland, A., Merisaari, H., Saunavaara, J., Parkkola, R., Lähdesmäki, T., Saukko, E., Nolvi, S., Kataja, E.-L., Korja, R., Karlsson, L., Karlsson, H., & Tuulari, J. J. (2022). Feasibility of FreeSurfer Processing for T1-Weighted Brain Images of 5-Year-Olds: Semiautomated Protocol of FinnBrain Neuroimaging Lab. *Frontiers in Neuroscience*, *16*. https://doi.org/10.3389/fnins.2022.874062
